# Metabolic Interactions Between Dynamic Bacterial Subpopulations

**DOI:** 10.1101/208686

**Authors:** Adam Z. Rosenthal, Yutao Qi, Sahand Hormoz, Jin Park, Sophia Hsin-Jung Li, Michael Elowitz

## Abstract

Within multi-species microbial communities, individual species are known to occupy distinct metabolic niches. By contrast, it has remained largely unclear whether and how metabolic specialization occurs within clonal bacterial populations. The possibility of such metabolic specialization in clonal populations raises several questions: Does specialization occur, and if it does, which metabolic processes are involved? How is specialization coordinated? How rapidly do cells switch between states? And finally, what functions might metabolic specialization provide? One potential function of metabolic specialization could be to manage overflow metabolites such as acetate, which presents a toxic challenge due to low pH, and protective pH-neutral overflow metabolites. Here we show that exponentially dividing Bacillus subtilis cultures divide into distinct interacting metabolic subpopulations including one population that produces acetate, and another population that differentially expresses metabolic genes for the production of acetoin, a pH-neutral storage molecule. These subpopulations grew at distinct rates, and cells switched dynamically between states, with acetate influencing the relative sizes of the different subpopulations. These results show that clonal populations can use metabolic specialization to control the environment through a process of dynamic, environmentally-sensitive state-switching.

## Introduction

The Gram-positive bacterium *Bacillus subtilis* has two preferred carbon sources: glucose and malate (Kleijn et al. 2010). When both of these carbon sources are available they are consumed simultaneously, generating growth rates that surpass those achieved with either substrate alone (Kleijn et al. 2010). Under conditions of rapid growth, co-consumption of glucose and malate leads to the accumulation of high levels of acetate (Kleijn et al. 2010). As a weak organic acid, acetate can be harmful to cells even in buffered medium (Rosenthal, Kim, and Gralla 2008). Acetate and related short-chain fatty acids enter the cell passively in the neutral form and then dissociate intracellularly, releasing a proton and transiently acidifying the cytoplasm (Russell and Diez-Gonzalez 1997; A. J. Roe et al. 1998). The intracellular dissociation of acetate also disrupts the cellular anion balance, with negative effects on metabolism (A. J. Roe et al. 1998; Andrew J. Roe et al. 2002) and transcription (Rosenthal, Kim, and Gralla 2008). When extracellular acetate levels rise to toxic levels the growing *Bacillus subtilis* culture consumes the acetate and produces acetoin, a non-toxic pH-neutral overflow metabolite that can be used as a carbon source in later growth stages (Speck and Freese 1973) (Fig. 1A).

**Figure 1:**
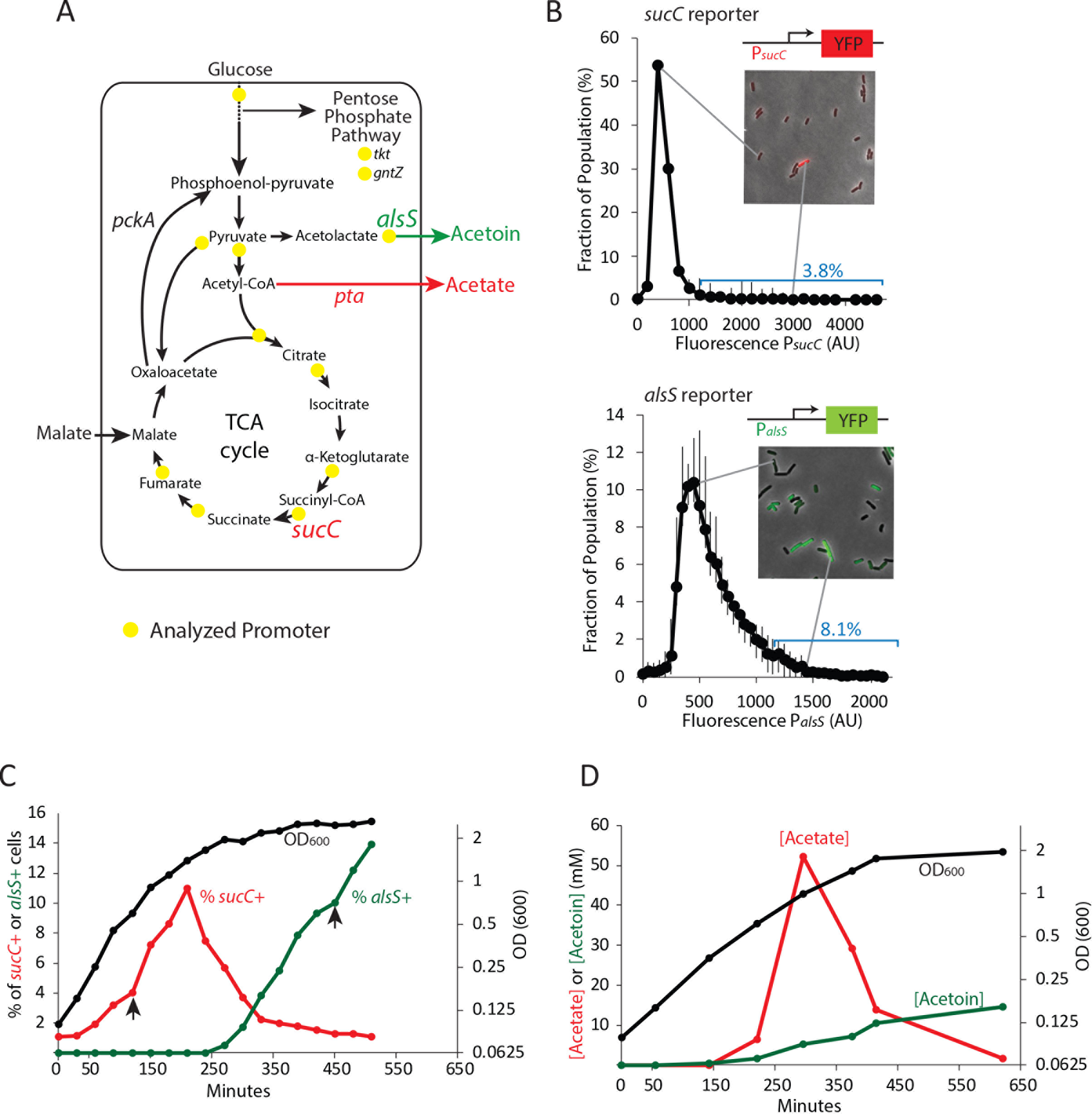
Two genes in central carbon metabolism are heterogeneously expressed in a clonal population of *B. subtilis*. **A)** *B. subtilis* uses glucose and malate as preferred carbon sources, and under aerobic culture conditions produces acetate and acetoin as major overflow metabolites. Promoter reporter strains were made for genes participating in the reactions marked with a yellow dot **B)** The heterogeneous expression of *sucC* (red line) and *alsS* (green line) is maximal at different timepoints along the growth curve (black line). Black arrows denote timepoints shown in figure 1C. **C)** Histograms depict the heterogeneous expression of the central metabolism genes *sucC* (left panel) and *alsS* (right panel). Insets using merged phase and fluorescence images show typical fields of cells, including cells in the high expressing tail of the distributions. **D)** A line graph depicting the accumulation of extracellular acetate and acetoin in the growth media during exponential and early stationary growth (OD_600_, black line). Acetate (red line) is released around mid-exponential phase, and is reabsorbed at a later time during which acetoin is produced (green line).

A biphasic growth strategy, in which acetate is produced to a toxic level and then reabsorbed and replaced by a non-toxic metabolite (Wolfe 2005), is common to many bacterial species and is important both for understanding the basic biology of bacterial growth in culture, and for applications in metabolic engineering (Papagianni 2012). However, it has generally been studied only at the population level, implicitly assuming a homogeneous progression of the entire culture from acetate producing to acetate detoxifying states. However, recent work at the single cell level suggests that bacterial systems can exhibit enormous heterogeneity in functional and gene expression states across diverse systems (Eldar et al. 2009a; Locke et al. 2011a; Süel et al. 2006; Levine et al. 2012). This prompts the questions of whether microbial cells differentiate into metabolically distinct subpopulations, and more specifically, whether acetate production and detoxification might occur in distinct cells specializing in acetate production or detoxification, respectively.

## Results

To address these questions we constructed a library of strains with reporters for key genes involved in central carbon metabolism, acetate production, and organic acid detoxification (Fig. 1A). We introduced a fluorescent protein (YFP) under the control of promoters for 13 different metabolic genes and stably incorporated them into the commonly used *sacA* site within the genome (Table S1), (Eldar et al. 2009a; Locke et al. 2011a). Using quantitative single-cell fluorescence microscopy, we analyzed the distribution of expression levels of these genes in individual cells at different times along the growth curve in buffered culture medium containing 0.4% glucose and 50 mM malate. To eliminate oxygen gradients, 10 mL cultures were grown in 250 mL flasks with rapid shaking (250 RPM). Most genes showed unimodal and relatively tight distributions (Figure S1) with coefficients of variation (CV)<25%). By contrast, expression of *sucC* and *alsS*, genes that encode succinate co-A ligase and acetolactate synthase, respectively, was more heterogeneous (Fig. 1B). We observed CVs greater than 30%, with 3.8% of P_*sucC*_-YFP and 8.1% of P_*alsS*_-YFP cells exhibiting high expression levels (≥2 standard deviations above the mean) at OD_600_ ~0.8 (*sucC*) and OD_600_ ~2 (*alsS*). Thus, at least two metabolic genes are heterogeneously expressed under these conditions.

To better understand when this heterogeneity emerges in batch culture, we performed a time course analysis of the fraction of *sucC* and *alsS* positive cells (≥2 standard deviations above the mean denoted *sucC+* and *alsS+*, Fig. 1C). We observed that the subpopulation of cells expressing *sucC* only exists transiently, in mid-to late-exponential phase (Fig. 1C). Furthermore, this dynamic pulse of *sucC+* cells coincided with the time and culture optical density at which acetate production was observed (~180-300 min, Fig. 1D). This observation suggested that *sucC* expression could be involved in acetate production. A parallel analysis of *alsS* expression revealed the opposite behavior, with *alsS* expression dynamics coinciding with a decrease in acetate and a concomitant increase in acetoin levels (Fig. 1C, D). This behavior is generally consistent with the known role of *alsS* in acetoin production in response to acetate toxicity (Speck and Freese 1973). Together, these results show that a dynamic change in acetate and acetoin levels in the culture overlaps with changes in the population fraction of *sucC* and *alsS* expressing cells.

A role for SucC in acetate production has not been studied previously. To understand the relationship between the *sucC+* subpopulation and acetate production, we used fluorescence activated cell sorting (FACS) of the P_*sucC*_YFP reporter strain to sort cells expressing YFP from a *SucC* promoter at the time of peak acetate levels, and performed RNAseq to compare gene expression profiles (Fig 2A, Fig. S2). As expected, *sucC* expression was elevated 2-fold in the *sucC+* sorted subpopulation (blue dot, Fig 2A). For most genes, we observed a broad correlation in gene expression between the two populations. However, RNAseq analysis with cuffdiff (Trapnell et al. 2010) and gene set enrichment analysis with GSEA (Subramanian et al. 2005) showed that genetic competence genes (Berka et al. 2002) were significantly enriched in the ~300 upregulated genes in the *sucC+* subpopulation (red dots and inset, Fig. 2A and Table S2; GSEA p <e-16). The *sucC+* population also exhibited increased expression of the phosphate acetyltransferase gene, *pta* (green dot, Fig. 2A), the enzyme that catalyzes the final step in overflow acetate production. *sucC+* cells apparently represent a distinct gene expression state that could be involved in acetate production.

**Figure 2:**
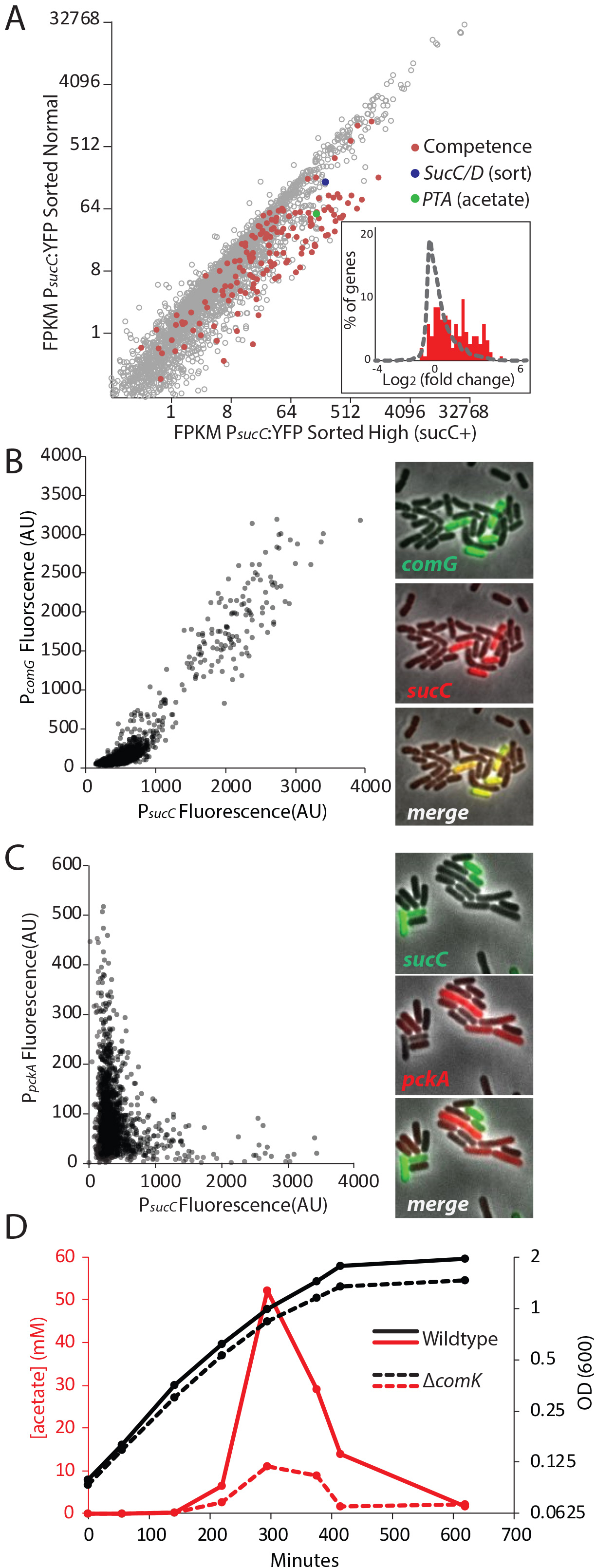
The heterogeneous expression of *sucC* is correlated with the genetic-competence regulon, and this metabolic state produces extracellular acetate. **A)** RNAseq of cells sorted at either a high or moderate *sucC* level reveals positive correlation between sucC expression, the competence program, and the acetate metabolism gene *pta*. Cells expressing YFP under the control of the *sucC* promoter were sorted at high or normal expression levels. Genes for genetic competence (red) and acetate production (*pta* - green) are higher in cells expressing high levels of *sucC*. The inset histogram shows a histogram of log_2_ fold change for all genes (grey) and the competence program genes (red) **B)** A scatter plot shows that the expression of *sucC* reporter is positively correlated with the *comG* reporter, a marker for the competence program. Each dot represents a single cell centered on the mean fluorescence of reporters for *sucC* and *comG*. The right panel shows fluorescent microscopy images taken from a typical field of cells **C)** A scatter plot shows that the expression of the *sucC* reporter is negatively correlated with expression of the metabolic gene *pckA*. Each dot represents a single cell centered on the mean fluorescence of reporters for *sucC* and *pckA*. Right panel shows fluorescent microscopy images taken from a typical field of cells. **D)** The competence gene expression program is necessary for the buildup of high levels of extracellular acetate. Growth curves demonstrate only a small difference in growth of wildtype strain (solid black line) or the competence-null Δ*comK* strain (dashed black line). However, maximal acetate buildup is approximately 5 fold higher in the wildtype strain (solid red line) than in a strain that is unable to produce the competent cell population (Δ*comK* dashed red lines).

Based on the strong correlation between *sucC* expression and competence gene expression in the RNAseq results (Fig. 2A), we next asked whether the *sucC+* subpopulation represented the competent state. To analyze the relationship between *sucC* expression and genetic competence in single cells, we constructed four dual reporter strains, expressing CFP from the *sucC* promoter and YFP from one of four competence promoters: *comG*, *comK*, *nucA* and *rapH* (Berka et al. 2002). Imaging revealed a clear positive correlation between *sucC* and the competence genes (Fig. 2B, Fig S3). This positive correlation was not general to all metabolic genes, as *sucC* expression was anti-correlated with *pckA* (fig 2C, Fig S4), a gene involved in phosphoenolpyruvate synthesis (Meyer and Stülke 2013). We note that *pta* and *sucC* were previously observed to be up-regulated in the competent state (Berka et al. 2002). Together, these results suggest that individual cells can exist in at least two distinct metabolic states, one of which represents genetically competent cells and involves increased expression of *sucC* and *pta*, among other genes.

We next assessed how competence might be linked to elevated acetate production. The competence system is controlled by a noise-excitable gene circuit that stochastically initiates transient episodes of differentiation in individual cells (Suel et al. 2007; Süel et al. 2006; Cağatay et al. 2009; Maamar, Raj, and Dubnau 2007;J. Hahn, Kong, and Dubnau 1994) To better understand the relationship between competence and acetate metabolism, we next asked whether activation of the competence system is necessary for increased *sucC* expression. Strains in which the competence master transcription factor *comK* is deleted (Table S1) exhibited reduced acetate production (Fig. 2D) and a loss of *sucC* as well as *comG* expression (Fig. S5). Although the competent state has been suggested to be involved in other functions, such as attachment, motility, antibiotic resistance, and DNA metabolism (Redfield 1993;Jeanette Hahn et al. 2015; Bakkali 2013; Finkel and Kolter 2001), a role in central carbon metabolism has not been appreciated. These results indicate that the *sucC* subpopulation is controlled by the competence system, linking competence both to an alternative metabolic state and to the control of acetate levels in culture.

**Figure 3:**
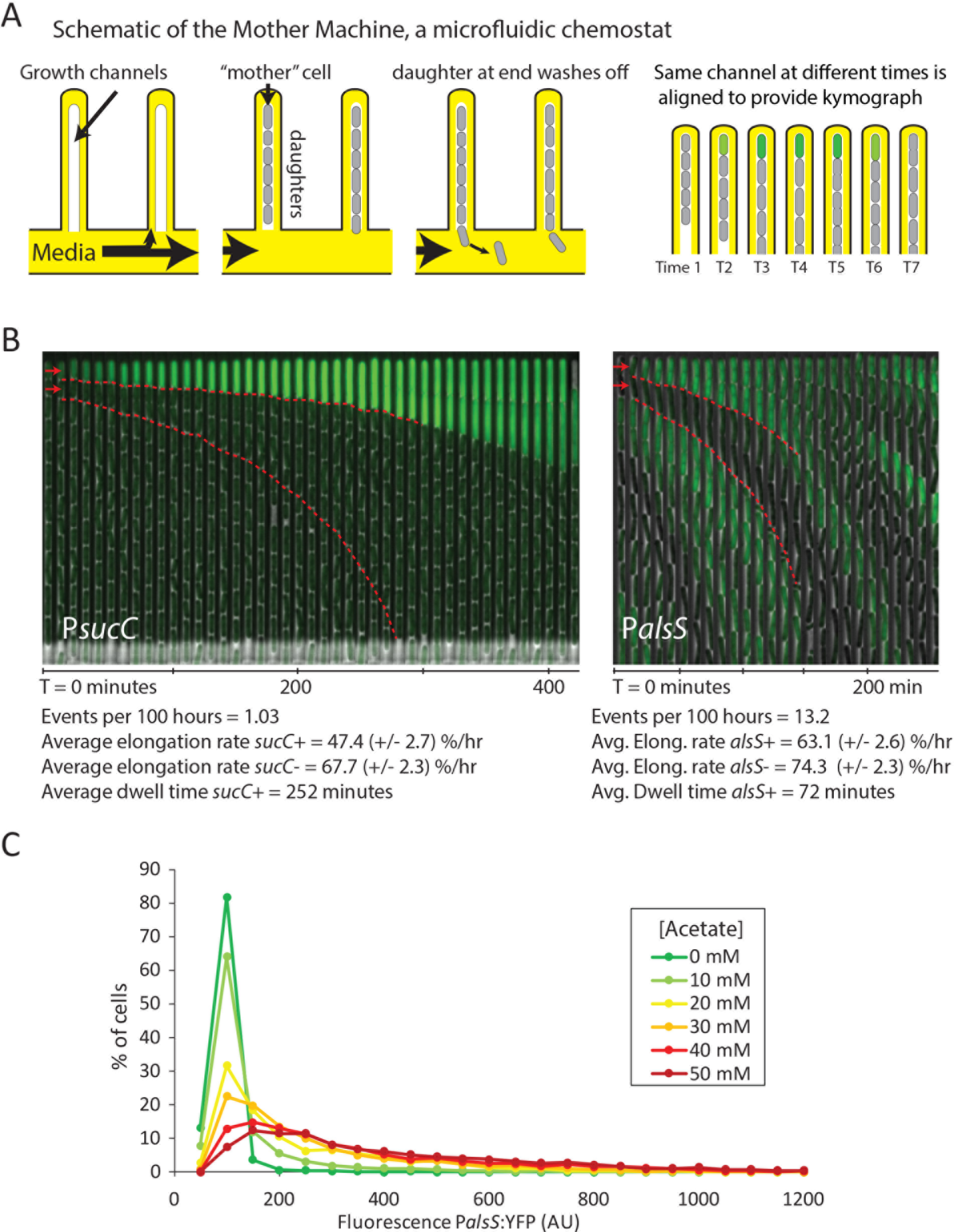
Cells switch in and out of the slower growing *sucC+* and *alsS+* states based on media conditions. **A)** A schematic of the Mother Machine microfluidic experiment. Cells are loaded into growth channels that are capped on one end and surrounded by flowing media. A “mother” cell settles at the capped end, and produces daughters. The daughters at the uncapped end of the growth channel are washed away by the current of media. Positions were filmed for up to 4 days, and for visualization purposes the images from each growth channel were cropped and aligned to generate a kymograph depicting time on the x-axis. **B)** Filmstrip kymographs from mothermachine experiments using conditioned media at OD_600_ 0.8 and *sucC* reporter strain (left panel) or conditioned media at OD_600_ 2.0 using *alsS* reporter strain (right panel). Dashed red lines show the trend of growth of two daughters, a *sucC+* and *sucC−* pair on the left panel and a *alsS+* and *alsS−* pair on the right panel. As seen from the slope of the trend lines and as indicated below the kymographs, the elongation rates of both *sucC+* and *alsS+* cells are slower than their counterparts. **C)** Extracellular acetate levels activate heterogeneous expression of the *alsS* promoter. A histogram shows the population of cells expressing different levels of YFP under control of the *alsS* promoter. When no acetate is added (green line), practically all cells are in the low expressing portion of the histogram. When extracellular acetate levels are added to mimic the maximal amount produced in the growth curve (orange and red lines) some cells remain in the low expressing portion of the histogram, but a correspondingly larger number of cells are in the long tail of high *alsS* expression.

**Figure 4:**
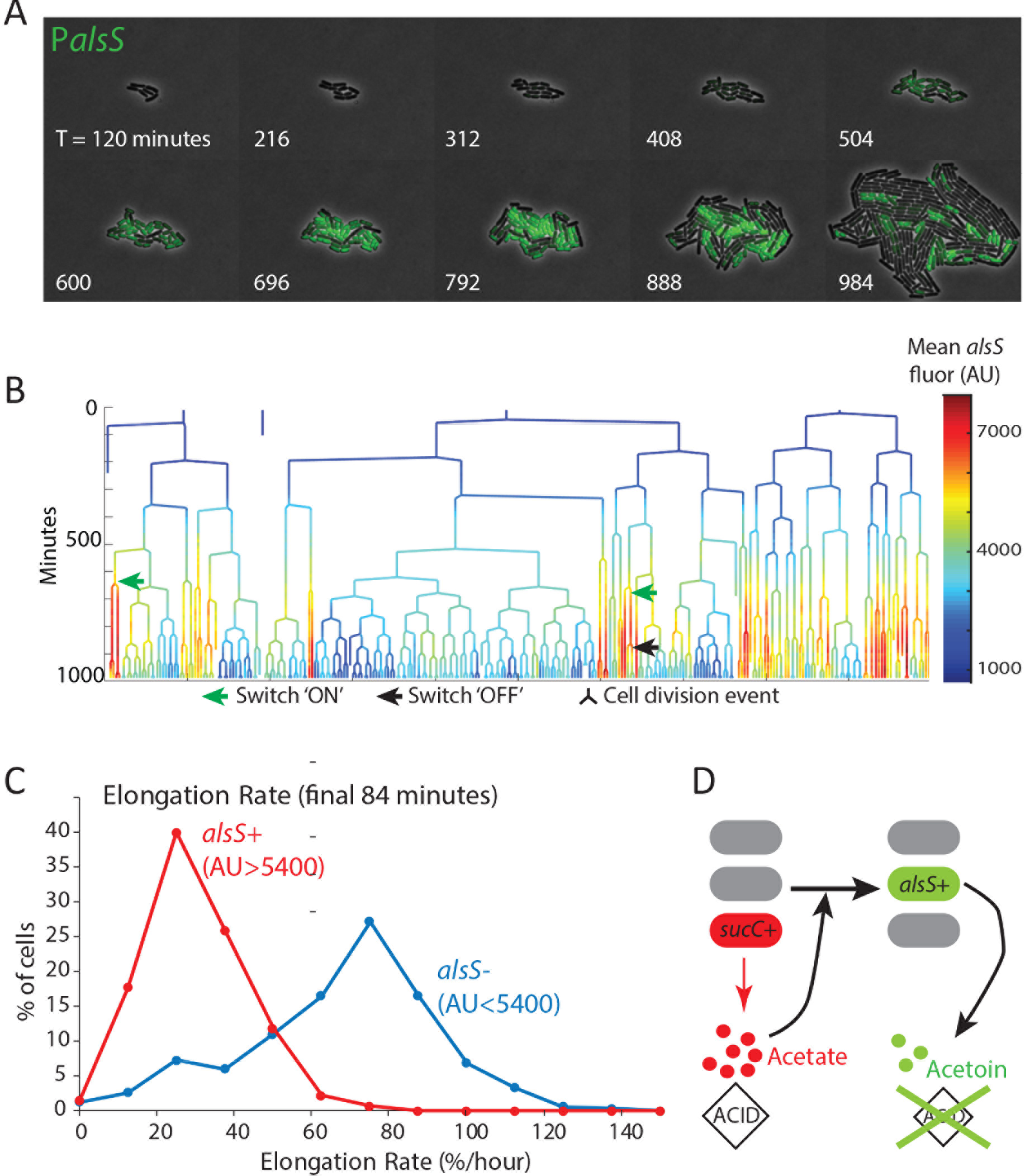
alsS+ cells have slower division and elongation rate than alsS-cells. **A)** A filmstrip of a representative timelapse experiment. Cells were grown on agarose pads containing acetate at a level that mimics mid-exponential phase. **B)** *AlsS+* cells divide more slowly. A genealogy tree depicts cell division events in the experiment shown in panel A. *alsS* levels are color coded by the heatmap on the right. Cells switch in and out of high *alsS* expression levels. Cells expressing high *alsS* levels (red and orange) divide more slowly than cells with low *alsS* levels (blue). **C)** *AlsS−* cells in the end of the experiment have faster elongation rates. Cells in the last 7 frames of the experiment which had arbitrary fluorescence levels greater than 5400 were designated *alsS+* and those expressing less were designated *alsS−*. The elongation rates of each group of cells was determined and plotted as a histogram. *alsS−* cells (blue line) had a median elongation rate of 65.2%/hr while *alsS+* cells (red line) had a median elongation rate of 19.96 %/hr **D)** A cartoon summary of the *sucC* and *alsS* interactions: In early growth stages a subset of cells become *sucC+*. These cells secrete acetate. As acetate levels build up they can become toxic. These high acetate levels activate some cells in the population to preferentially express metabolic genes for the production of acetoin. Acetoin, a non-toxic pH-neutral metabolite, replaces acetate in the media.

To better understand the dynamics with which cell switch into the competence and later into the *alsS+*, we used the “Mother Machine” microfluidic device (Fig 4A), to conduct long-term analysis of individual cells over tens of cell generations under chemostatic conditions (Wang et al. 2010; Norman et al. 2013). We set up the Mother Machine as described previously (Norman et al. 2013), but cultured cells with conditioned media obtained from batch growth of *B. subtilis* cultures at different final optical densities. Specifically, we used media from cultures at OD_600_ 0.8 and OD_600_ 2.0, points during the peak of *sucC+* or *alsS+* expression, respectively. This approach provides the simplicity of long-term chemostatic analysis with the ability to compare cellular behavior at different culture time-points.

Using the Mother Machine, we analyzed cell lineages for up to 4 days (approximately 60 generations) for a total of 1,400 cell generations. With *sucC*-inducing media (conditioned at OD 0.8), we observed rare episodes of *sucC* activation in some cells, lasting for approximately four hours each (252 ± 89 minutes). Consistent with previous analysis of competence dynamics (Süel et al. 2006), these cells divided infrequently, and grew more slowly than other cells in the same movies (elongation rates of 47.4 ± 2.7 %/hr and 67.7 ± 2.3 %/hr, respectively). Cells in the activated state could switch out of the *sucC+* state and resume normal growth rates (Fig. S6, Movie S1). Under these conditions, we did not observe activation of *alsS* expression. By contrast, in the OD 2.0 conditioned media we did not observe activation of *sucC* expression, but we did observe frequent pulses of *alsS* gene expression. *alsS+* cells grew at a slightly reduced elongation rate (63% ± 2.6 %/hr increase compared to 74% ± 2.3 %/hr for alsS- cells, Fig. 3B, right panel, Movie S2). Together, these results provide the rates of transitions into the *sucC+* (competent) and *alsS+* gene expression states, and show that these states have altered growth rates and respond to medium composition.

These results suggested the possibility that acetate predominantly produced by *sucC+* cells early in the growth could be inducing cell switching to the *alsS+* state in later growth stages, when it accumulates to toxic levels. However, many components could differ between the OD_600_ 0.8 and OD_600_ 2.0 medium. To determine whether acetate was sufficient to affect *alsS* expression, we cultured reporter cells in varying levels of acetate, in unconditioned liquid medium, and quantified the fraction of *alsS+* cells. We observed both a systematic increase in the distribution of alsS expression levels, and in the fraction of cells in the high expressing “tail” of the distribution (Fig. 3C).

The Mother Machine is ideal for analyzing cells over multiple generations in a relatively constant environment but not ideal for analyzing responses to environmental changes that happen as a consequence of growth. To analyze more dynamic environmental conditions, we designed microcolony pad experiments in which acetate was added to standard microcolony medium (Eldar et al. 2009a; Locke et al. 2011a; Young et al. 2011a) to 20mM, the acetate concentration present in mid-exponential phase (Figure 1B) (Speck and Freese 1973). In these experiments (Fig. 4), all cells started with a low growth rate, likely owing to the initial acetate present in the growth media. As cells divided, a subpopulation of approximately half of cells switched on high levels of *alsS* expression within 7 to 10 hours (Fig. 4, Fig. S6, Movies S3-5). As growth progressed, these *alsS+* cells exhibited a reduced growth rate, similar to that of the original culture. However, a distinct subpopulation with approximately 2.5-fold lower *alsS* expression emerged, becoming greater than 70% of the population. These cells exhibited a faster division rate (Fig 4B, Fig s7) and faster elongation rate (Fig 4C, Fig S8).

The fast growing *alsS−* cells that appeared late in pad growth experiments had a large growth advantage compared to the slow growing *alsS+* cells (median elongation rates of 65 %/hr and 20%/hr, respectively). In general, growth in the chemostatic Mother Machine using conditioned media is faster than on a pad with non-conditioned media containing acetate. However, the difference between *alsS-* and *alsS+* cells is much smaller in microfluidic growth (74%/hr vs 63%/hr for *alsS-* and *alsS+*). This finding is consistent with the established role of acetoin as a molecule secreted to counter the pH and anion producing toxic effect of secreted short chain fatty acids,including acetate (Xiao and Xu 2007; Speck and Freese 1973). In the pad environment, transient activation of genes for the production of the pH-protective acetoin in the *alsS+* cells has the potential to produce a milder growth environment which may enable other cells to grow faster (Fig 4D). By contrast, in the mother-machine experiments (figure 3), while cells switch in and out of the *alsS* metabolic state, the chemostatic nature of the device minimizes their ability to impact the growth rates of their neighbors.

## Discussion

The accumulation and reabsorption of acetate is a classic hallmark of bacterial growth in aerobic conditions, common to many bacteria including *B. subtilis* and *E. coli* (Wolfe 2005). This regime, commonly referred to as the “acetate switch” allows for very rapid growth when acetate is produced. When acetate levels and associated acidity reach toxic levels, acetate is reabsorbed and replaced with pH-neutral overflow metabolites such as acetoin (Wolfe 2005; Speck and Freese 1973). Growth strategies in which a preferred toxic overflow metabolite is produced under aerobic conditions are also used by other organisms that expel and control different fermented toxic overflow metabolites, including ethanol fermentation by yeast (Otterstedt et al. 2004) and lactic acid in lactobacillus species (Borch and Molin 1989). Interestingly, in the fermentation of ethanol by the budding yeast *Saccharomyces cerevisiae*, ethanol is produced in dynamic bursts in which some cells switch in and out of fermentative metabolism. These bursts in yeast can be synchronized in chemostat growth (Tu et al. 2005), but also appear in batch culture (Silverman et al. 2010), in which different cell subpopulations express fermentative or respiratory genes. In yeast, the single cell dynamics, mechanisms, and role of these bursts have not been fully elucidated. However, the presence of metabolically specialized subpopulations of cells in both eukaryotes and bacteria suggests that segregating different fermentative or respiratory pathways into individual cells may be a general strategy. Such a separation of metabolic activities in different subpopulations may be useful in avoiding metabolic incompatibilities (Brandriss and Magasanik 1981; Ackermann 2015; Kumar, Mella-Herrera, and Golden 2010), controlling cellular challenges such as reducing potential (Liu et al. 2017), or for optimizing enzyme and substrate scaling, where locally high concentrations may be needed for efficient enzymatic conversion to occur (Nikel et al. 2014; Ackermann 2015). Better understanding of the principles that govern such metabolic activity segregation will have implications for efforts to design robust metabolic engineering and industrial fermentation approaches. This is especially true for commonly used industrial strains which naturally produce multiple fermentation products. For example, *E. coli* strains simultaneously produce five different fermentation products during mixed-acid fermentation (Clark 1989).

The rapid dumping of toxic overflow products also has a role in the context of competition within a multi-species environment. In such environments, a quick buildup of toxic products can be advantageous to ward off competing species. In the case of human infectious disease, the buildup of byproducts such as lactic acid from normal flora species limits infection by pathogens that are not lactic acid specialists (O′Hanlon, Moench, and Cone 2013). Likewise, in industrial fermenters and microbial food fermentation secreted overflow metabolites including acetate and ethanol limit contamination. Additionally, if a particular metabolic niche is transient, as in the case of acetate in *B. subtilis* colony growth or batch culture, a strategy in which cells can switch in and out of metabolic states can be advantageous to an alternative scenario in which multiple strains are evolutionarily “locked” into distinct specialist roles. This is especially true if the metabolic niche (e.g. acetate) is short lived, because a “locked” specialist strain would greatly diminish in numbers and even risk extinction during periods of growth when acetate is absent, and would subsequently be in diminished numbers once conditions are favorable.

In this study, we link the presence of extracellular acetate with the activation of *alsS* in a subset of cells. In the case of competence and *sucC*, which are both controlled by the master regulator *comK*, a large literature describes the roles of quorum sensing in the activation of the competence program (D. Dubnau 1991). For maximal competence to manifest specific growth requirements and media components are needed (David Dubnau 1982), raising the possibility that alongside quorum sensing peptides, secreted metabolic byproducts also play a role in this process.

Going beyond microbial systems, cell-cell heterogeneity can be advantageous as a ‘bet-hedging’ strategy both for microbial and cancer cells (Veening, Smits, and Kuipers 2008). In such cases, cell-cell heterogeneity enables the population as a whole to withstand unforeseen challenges, such as antibiotic or chemotherapeutic drugs (Sharma et al. 2010; Rotem et al. 2010), or metabolic shifts (Solopova et al. 2014). By contrast to a simple bet-hedge, the emergence of *alsS+* acetoin producing populations described here arise as a response to an anticipated challenge that is part of the growth progression in conditions favoring weak organic acid production. Thus, unlike in bet-hedging, metabolic state switching provides a predictable benefit in a more deterministic environmental setting.

## Materials and Methods

### Plasmid design

Plasmids for the integration of fluorescent reporters were made as previously reported (Eldar et al. 2009b). YFP promoter reporters were cloned into the ECE174 backbone plasmid which uses sacA integration site and encodes chloramphenicol resistance (R. Middleton, obtained from the Bacillus Genetic Stock Center). CFP promoter reporters were cloned into the pDL30 backbone which uses amyE integration sites and encodes spectinomycin resistance (obtained from the Bacillus Genetic Stock Center). A constitutive RFP reporter, using a minimal sigA promoter, was used for image segmentation as previously reported (Locke et al. 2011b). pAZR1: (Pcggr:alsS/D) is a plasmid for the integration of a constitutive promoter (cggr promoter) to drive the constitutive expression of the alsS/D regulon from its native site. The plasmid was constructed by Gibson cloning (Gibson et al. 2009). A markerless deletion of alsS/D was made using the alsS/D strain of the BKE collection and the pdr244 plasmid, both obtained from the BGSC, followed by selection.

### Bacterial strains

All strains were made by genomic integration into the genome. Fluorescent reporters were integrated into either the *sacA* (YFP) or the *amyE* (CFP) loci as previously described (Locke et al. 2011b). A constitutive RFP color was utilized, relying on constitutive expression of a partial ptrpE promoter reporter driving mCherry expression, which was inserted into the ppsB locus as previously (Locke et al. 2011b). Non chaining strains for microfluidic mother-machine experiments used a lytF overexpression construct as previously reported. Strain information is included in the supplementary table.

### Growth Conditions

Strains were started from glycerol stocks and grown in M9 minimal media prepared according to the directions of the manufacturer (BD - difco, Franklin Lakes NJ). Base media was supplemented with 0.4% glucose and a cocktail of trace metals (Leadbetter et al. 1999) Malate was added to growing cultures at OD 0.4-0.5 as per previous publications (Buescher et al. 2012; Kleijn et al. 2009). Samples for fluorescence microscopy were prepared using agarose pads for either snapshot analysis (timepoint measurements) or pad movies, as previously described by our laboratory (Young et al. 2011b).

### Microscopy

Images were acquired using a Nikon inverted TI-E microscope via a coolsnap HQ2 camera. Commercially available software (Metamorph) controlled the stage, microscope, camera, and shutters. Fluorescent illumination was provided by a Sola Light Engine LED source (Lumencor). Temperature was kept at 37°C using an enclosed microscope chamber (Nikon) attached to a temperature sensitive heat exchanger. All experiments used a Phase 100x Plan Apo (NA 1.4) objective. Filter sets used were Chroma #41027 (mCh), Chroma #41028 (YFP), and Chroma #31044v2 (CFP).

### Measurements of secreted acetate and acetoin

Media was collected from growing cultures by centrifuging 500 ul culture samples at 5000 g for 2 minutes and filtering the supernatant in 0.2uM syringe filters. Clarified conditioned media samples were placed into glass sample vials and run at the Caltech environmental analysis center using an Aminex HPX-87H column (Bio-Rad, Rockville NY) in an Agilent 1100 HPLC with UV and Refractive Index detectors with elution using 0.013 N H_2_SO_4_ at ambient temperature and 36 ml/hour flow as described in (Leadbetter et al. 1999). Standards of acetate and acetoin were prepared in uninoculated growth media, and diluted to produce a standard curve.

### Microfuidic Mothermachine experiments

Microfluidic experiments used the mothermachine devices described in (REFS Wang Jun 2010, Norman Losick 2013). SU80 wafers were made based on masks provided by the Losick lab. PDMS devices were prepared by pouring degassed Sylgard 184 PDMS silicone (corning corporation, Corning NY) onto wafers and curing the molds for a minimum of 8 hours at 65°C. Cured PDMS devices were bonded onto microscopy coverslips (60×22 mm, Gold Seal coverslips Thermo Fisher Scientific, Waltham MA) by plasma cleaning. Plasma bonding was done in a PDC32G plasma cleaner (Harrick Plasma Ithaca, NY) set to chamber pressure between 600-700 microns. Coverslips were cleaned separately for 1 minute, and then the devices and coverslips were cleaned jointly for 20 seconds. Device bonding was immediately done by inverting the plasma treated device onto the treated coverslips. After bonding the devices were cured for an additional 4 hours at 65°C. Devices were kept for up to 2 weeks in the dark at room temperature. Before use, holes for inlet and outlet were punched using a biopsy punch and each device was passivated by loading the channels using growth-media containing 1 mg/ml BSA using a handheld micro-pippete and a 20uM tip. Cells were loaded by flowing a concentrated cell culture (OD 2.0) and letting cells reach the growth chambers by waiting for 30 minutes. Devices were placed on an inverted Nikon TiE microscope and growth media was flowed using syringe pumps set to a flow rate of 50-100 ul per hour. Media used in microfluidic mother machine experiments was conditioned media taken from batch growth cultures. Media for the sucC/competence experiments contained media conditioned by growth on glucose/malate media until OD 0.8. Conditioned media used for alsS movies was from OD 2.0. Fluorescent Images were captured using a CoolSnap HQ2 and analyzed with custom MATLAB software and in imageJ.

Cultures of cells expressing YFP under the control of *sucC* (strain AZRE1) were grown in M9 glucose-malate media. Cells in mid log phase (OD 0.8) were fixed in 4% formaldehyde for 10 minutes at room temperature. Fixed cells were washed twice in Tris pH 7, and gently filtered using a 5.0 uM filter to remove clumps and chains. Cells were sorted on either a MoFlo astrios cell sorter or a BSfacsARIA in the USC medical school sorting facility. Cells sorted for either high YFP fluorescence or regular fluorescence (a minimum of 200,000 cells) were collected into tubes containing RNA protect (Qiagen, Hilden, Germany). Sorted samples were centrifuged and cells were rehydrated in 240ul qiagen PKD buffer (FFPE miRNEASY kit, Qiagen). Cells were lysed by the addition of 10ul lysosome solution for 10 minutes followed by bead beating for 2 minutes in high setting. Samples were further processed using the qiagen FFPE miRNA kit. Libraries were prepared using the Epicentre Scriptseq V2 kit, follow the directions for highly fragmented DNA. Libraries were sequenced at the Caltech Millard and Muriel Jacobs sequencing facility. Analysis followed the standard Galaxy RNAseq workflow (grooming, trimming, bowtie mapping, and cuff-diff and cuff-links) (Afgan et al. 2016).

### Agarose pad timelapse experiments

Agarose pad experiments were done as previously described (Young et al. 2011b), with the following exceptions: Cells were spotted on agarose pads made with standard pad movie media (Young et al. 2011b) which is a Spizizen’s minimal media with 0.4% glucose to which acetate was added to a final concentration of 20mM, to mimic acetate concentrations at mid exponential phase. Cells were allowed to acclimate to the agarose pad growth condition for ~2 h, before the start of imaging. Images were acquired from multiple fields every 12 minutes for a total of 22 hours. Movie analysis was performed in Matlab using the Schnitzcells analysis package (Young et al. 2011b) with slight edits. The current version of this analysis package is available at http://www.elowitz.caltech.edu

## Acknowledgements

We thank Jared Leadbetter, Ned Wingreen, Xinning Zhang, Avigdor Eldar, Joe Levine, Eric Matson, and members of the Elowitz lab for discussions and comments. This research was supported by the by Defense Advanced Research Projects Agency Biochronicity Grant DARPA-BAA-11-66, NIH R01GM079771, and a Caltech CEMI (Center for Environmental Microbial Interactions at Caltech Interactions) grant (AZR).

## Supplementary Figure, Table, and Movie Legends

**Supplementary Figure 1:** Histograms of metabolic genes with low cell to cell heterogeneity. The expression level of promoter reporters for the genes *acoA*, *citB*, *citZ*, *fumC*, *gntZ*, *odhA*, *pdhA*, *ptsG*, *pycA*. *sdhC*, *and tkt* in individual cells are shown. Cells with similar expression levels were binned and values for each bin are displayed in the histograms. Cells were collected from cultures grown in M9 Glucose/Malate media.

**Supplementary Figure 2:** FACS sorting parameters for sorting *sucC+* and normal cells for RNAseq experiments. Cells carrying a promoter reporter expressing YFP under control of the *sucC* promoter were sorted based on relative YFP fluorescence. The G3 sorting gate (blue regions corresponding to 4.3% of analyzed events) was designated “P*sucC* High”. The G2 sorting gate (green region corresponding to 88.45% of analyzed events) was designated “P*sucC* Normal”

**Supplementary Figure 3:** Competence genes are positively correlated with the TCA cycle gene *sucC*.

Scatter plots show that the expression of *sucC* reporter (CFP) is positively correlated with expression of the YFP promoter reporters for the classical competence program genes *comK*, *nucA*, and *rapH* in single cells. Each dot represents a single cell centered based on the mean fluorescence of fluorescent reporters for *sucC* and the competence gene.

**Supplementary Figure 4:** Competence genes are negatively correlated with the metabolic gene *pckA*

Three replicate scatter plots show that the expression of *sucC* reporter (CFP) is negatively correlated with expression of the YFP promoter reporter for the Phosphoenolpyruvate metabolism gene *pckA* in single cells. Each dot represents a single cell centered based on the mean fluorescence of fluorescent reporters for *sucC* and the competence gene. The red dashed lines show regions used in the figure to test for anticorrelation using the Fisher Exact Test. P values for the Fisher exact test (anticorrelation) are displayed for each replicate, as well as R values.

**Supplementary Figure 5:** The heterogeneous expression of *sucC* and *comG* are dependent on the competence master regulator comK. The expression level of promoter reporters for the genes *sucC* (CFP), *and comG* (YFP) in individual cells were determined in wildtype cells (blue dots and blue shading) as well as in cells in which the master competence regulator comK was deleted (ΔcomK) (red dots and red shading). Cells with similar expression levels were binned and values for each bin are displayed in the histograms. Cells were collected from cultures grown in M9 Glucose/Malate media. The wildtype cultures had greater than 4% of cells expressing the reporter at levels >2 standard deviations above the mean. Values in this tail of the distribution are showed in the zoom-in windows below the main panels.

**Supplementary Figure 6:**Filmstrip kymographs from mother-machine experiments using conditioned media at OD_600_ 0.8 and *sucC* reporter strain

**Supplementary Figure 7:** Cells expressing high levels of alsS divide more slowly than cells expressing lower levels. The genealogy plots from three replicate pad experiments are provided. The movies from each movie are included as supplementary pad movies. Time in the movie is indicated on the tree ordinate (Y axis). The level of alsS is indicated by the provided heatmap.

**Supplementary Figure 8:** Cells expressing high levels of alsS elongate more slowly than cells expressing lower levels in the end of the pad culture experiment. **A)** The elongation rate of each cell throughout the entire length of the experiment is provided in the ordinate (Y axis) for all different times in the movie. The level of *alsS* promoter reporter fluorescence is indicated in the provided heatmap. Early in the movie all cells have low PalsS expression levels and slow elongation rate. Near the end of the movie cells expressing high levels of *alsS* (yellow, orange, and red dots) appear, having a slow elongation rate. Additionally, a population of low expressing cells (blue) with a faster elongation rate also appear near the end of the experiment. **B)** The level of alsS in the final few frames is bimodal. Cells from the final frames were binned according to the level of alsS fluorescence. A low expressing population (AU<5400) and a high expressing population (AU>5400) are present. **C)** The elongation rate in the final frames is higher for cells expressing low levels of alsS. Cells were designated high and low expressing based on the parameters in panel (B). Cells from each group (alsS high and alsS low) were binned based on elongation rates and bins were displayed as a histogram. The distribution of growth rates from cells expressing high *alsS* levels (red line) is left shifted (lower elongation values) compared to that of cells expressing high *alsS* levels (blue line).

**Supplementary Table 1:** List of strains used in this work.

**Supplementary Table 2:** Genes significantly differentially regulated in cells sorted by P_*alsS*_:YFP fluorescence.

Genes that are significantly differentially regulated in the sorting and RNAseq experiments are sorted based on P-values. Genes in green text have P-values of 5.0 e-5 or smaller

**Supplementary Table 3:** Genes that were not significantly regulated in cells sorted by P_*alsS*_:YFP fluorescence.

## References

Ackermann, Martin. 2015. “A Functional Perspective on Phenotypic Heterogeneity in Microorganisms.” Nature Reviews. Microbiology 13 (8): 497–508.

Afgan, Enis, Dannon Baker, Marius van den Beek, Daniel Blankenberg, Dave Bouvier, Martin Čech, John Chilton, et al. 2016. “The Galaxy Platform for Accessible, Reproducible and Collaborative Biomedical Analyses: 2016 Update.” Nucleic Acids Research 44 (W1): W3–10.

Bakkali, M. 2013. “Could DNA Uptake Be a Side Effect of Bacterial Adhesion and Twitching Motility?” Archives of Microbiology 195 (4): 279–89.

Berka, Randy M., Jeanette Hahn, Mark Albano, Irena Draskovic, Marjan Persuh, Xianju Cui, Alan Sloma, William Widner, and David Dubnau. 2002. “Microarray Analysis of the Bacillus Subtilis K-State: Genome-Wide Expression Changes Dependent on ComK.” Molecular Microbiology 43 (5): 1331–45.

Borch, Elisabeth, and Goran Molin. 1989. “The Aerobic Growth and Product Formation of Lactobacillus, Leuconostoc, Brochothrix, and Carnobacterium in Batch Cultures.” Applied Microbiology and Biotechnology 30 (1). doi:10.1007/bf00256001

Brandriss, M. C., and B. Magasanik. 1981. “Subcellular Compartmentation in Control of Converging Pathways for Proline and Arginine Metabolism in Saccharomyces Cerevisiae.” Journal of Bacteriology 145 (3): 1359–64.

Buescher, Joerg Martin, Wolfram Liebermeister, Matthieu Jules, Markus Uhr, Jan Muntel, Eric Botella, Bernd Hessling, et al. 2012. “Global Network Reorganization during Dynamic Adaptations of Bacillus Subtilis Metabolism.” Science 335 (6072): 1099–1103.

Cağatay, Tolga, Marc Turcotte, Michael B. Elowitz, Jordi Garcia-Ojalvo, and Gürol M. Süel. 2009. “Architecture-Dependent Noise Discriminates Functionally Analogous Differentiation Circuits.” Cell 139 (3): 512–22.

Clark, D. 1989. “The Fermentation Pathways of Escherichia Coli.” FEMS Microbiology Letters 63(3): 223–34.

Dubnau, D. 1991. “The Regulation of Genetic Competence in Bacillus Subtilis.” Molecular Microbiology 5 (1): 11–18.

Dubnau, David. 1982. “Genetic Transformation in Bacillus Subtilis.” In Bacillus Subtilis, 147–78.

Eldar, Avigdor, Vasant K. Chary, Panagiotis Xenopoulos, Michelle E. Fontes, Oliver C. Losón,Jonathan Dworkin, Patrick J. Piggot, and Michael B. Elowitz. 2009a. “Partial Penetrance Facilitates Developmental Evolution in Bacteria.” Nature 460 (7254): 510–14.

Finkel, S. E., and R. Kolter. 2001. “DNA as a Nutrient: Novel Role for Bacterial Competence Gene Homologs.” Journal of Bacteriology 183 (21): 6288–93.

Gibson, Daniel G., Lei Young, Ray-Yuan Chuang, J. Craig Venter, Clyde A. Hutchison3rd, and Hamilton O. Smith. 2009. “Enzymatic Assembly of DNA Molecules up to Several Hundred Kilobases.” Nature Methods 6 (5): 343–45.

Hahn, Jeanette, Andrew W. Tanner, Valerie J. Carabetta, Ileana M. Cristea, and David Dubnau. 2015. “ComGA-RelA Interaction and Persistence in the Bacillus Subtilis K-State.” Molecular Microbiology 97 (3): 454–71.

Hahn, J., L. Kong, and D. Dubnau. 1994. “The Regulation of Competence Transcription Factor Synthesis Constitutes a Critical Control Point in the Regulation of Competence in Bacillus Subtilis”. Journal of Bacteriology 176 (18): 5753–61.

Kleijn, Roelco J., Joerg M. Buescher, Ludovic Le Chat, Matthieu Jules, Stephane Aymerich, and Uwe Sauer. 2009. “Metabolic Fluxes during Strong Carbon Catabolite Repression by Malate inBacillus Subtilis.” The Journal of Biological Chemistry 285 (3): 1587–96.

Kumar, Krithika, Rodrigo A. Mella-Herrera, and James W. Golden. 2010. “Cyanobacterial Heterocysts.” Cold Spring Harbor Perspectives in Biology 2 (4): a000315.

Leadbetter, J. R., T. M. Schmidt, J. R. Graber, and J. A. Breznak. 1999. “Acetogenesis from H2 plus CO2 by Spirochetes from Termite Guts.” Science 283 (5402): 686–89.

Levine, Joe H., Michelle E. Fontes, Jonathan Dworkin, and Michael B. Elowitz. 2012. “Pulsed Feedback Defers Cellular Differentiation.” PLoS Biology 10 (1): e1001252.

Liu, Jintao, Rosa Martinez-Corral, Arthur Prindle, Dong-Yeon D. Lee, Joseph Larkin, Marçal Gabalda-Sagarra, Jordi Garcia-Ojalvo, and Gürol M. Süel. 2017. “Coupling between Distant Biofilms and Emergence of Nutrient Time-Sharing.” Science 356 (6338): 638–42.

Locke, James C. W., Jonathan W. Young, Michelle Fontes, María Jesús Hernández Jiménez, and Michael B. Elowitz. 2011a. “Stochastic Pulse Regulation in Bacterial Stress Response.” Science 334 (6054): 366–69.

Maamar, Hédia, Arjun Raj, and David Dubnau. 2007. “Noise in Gene Expression Determines Cell Fate in Bacillus Subtilis.” Science 317 (5837): 526–29.

Meyer, Frederik M., and Jörg Stülke. 2013. “Malate Metabolism in Bacillus Subtilis: Distinct Roles for Three Classes of Malate-Oxidizing Enzymes.” FEMS Microbiology Letters 339 (1): 17–22.

Nikel, Pablo Iván, Rafael Silva-Rocha, Ilaria Benedetti, and Víctor de Lorenzo. 2014. “The Private Life of Environmental Bacteria: Pollutant Biodegradation at the Single Cell Level.” Environmental Microbiology 16 (3): 628–42.

Norman, Thomas M., Nathan D. Lord, Johan Paulsson, and Richard Losick. 2013. “Memory and Modularity in Cell-Fate Decision Making.” Nature 503 (7477): 481–86.

O’Hanlon, Deirdre E., Thomas R. Moench, and Richard A. Cone. 2013. “Vaginal pH and “Microbicidal Lactic Acid When Lactobacilli Dominate the Microbiota.” PloS One 8 (11): e80074.

Otterstedt, Karin, Christer Larsson, Roslyn M. Bill, Anders Ståhlberg, Eckhard Boles, Stefan Hohmann, and Lena Gustafsson. 2004. “Switching the Mode of Metabolism in the Yeast Saccharomyces Cerevisiae.” EMBO Reports 5 (5): 532–37.

Papagianni, Maria. 2012. “Recent Advances in Engineering the Central Carbon Metabolism of Industrially Important Bacteria”. Microbial Cell Factories 11 (April): 50.

Redfield, R. J. 1993. “Genes for Breakfast: The Have-Your-Cake-and-Eat-It-Too of Bacterial Transformation.” The Journal of Heredity 84 (5): 400–404.

Roe, A. J., D. McLaggan, I. Davidson, C. O’Byrne, and I. R. Booth. 1998. “Perturbation of Anion Balance during Inhibition of Growth of Escherichia Coli by Weak Acids.” Journal of Bacteriology 180 (4): 767–72.

Roe, Andrew J., Conor O’Byrne, Debra McLaggan, and Ian R. Booth. 2002. “Inhibition of Escherichia Coli Growth by Acetic Acid: A Problem with Methionine Biosynthesis and Homocysteine Toxicity.” Microbiology 148 (Pt 7): 2215–22.

Rosenthal, Adam Z., Youngbae Kim, and Jay D. Gralla. 2008. “Regulation of Transcription by Acetate in Escherichia Coli: In Vivo and in Vitro Comparisons.”Molecular Microbiology 68(4):907–17.

Rotem Eitan, Adiel Loinger, Irine Ronin, Irit Levin-Reisman, Chana Gabay, Noam Shoresh, Ofer Biham, and Nathalie Q. Balaban. 2010. “Regulation of Phenotypic Variability by a Threshold-Based Mechanism Underlies Bacterial Persistence.” Proceedings of the National Academy of Sciences of the United States of America 107 (28): 12541–46.

Russell, James B., and Francisco Diez-Gonzalez. 1997. “The Effects of Fermentation Acids on Bacterial Growth.” In Advances in Microbial Physiology, 205–34.

Sharma, Sreenath V., Diana Y. Lee, Bihua Li, Margaret P. Quinlan, Fumiyuki Takahashi, Shyamala Maheswaran, Ultan McDermott, et al. 2010. “A Chromatin-Mediated Reversible Drug-Tolerant State in Cancer Cell Subpopulations.” Cell 141 (1): 69–80.

Silverman, Sanford J., Allegra A. Petti, Nikolai Slavov, Lance Parsons, Ryan Briehof, Stephan Y. Thiberge, Daniel Zenklusen, et al. 2010. “Metabolic Cycling in Single Yeast Cells from Unsynchronized Steady-State Populations Limited on Glucose or Phosphate.” Proceedings of the National Academy of Sciences of the United States of America 107 (15): 6946–51.

Solopova, Ana, Jordi van Gestel, Franz J. Weissing, Herwig Bachmann, Bas Teusink, Jan Kok, and Oscar P. Kuipers. 2014. “Bet-Hedging during Bacterial Diauxic Shift.” Proceedings of the National Academy of Sciences of the United States of America 111 (20): 7427–32.

Speck, E. L., and E. Freese. 1973. “Control of Metabolite Secretion in Bacillus Subtilis.” Journal of General Microbiology 78 (2): 261–75.

Subramanian, Aravind, Pablo Tamayo, Vamsi K. Mootha, Sayan Mukherjee, Benjamin L. Ebert, Michael A. Gillette, Amanda Paulovich, et al. 2005. “Gene Set Enrichment Analysis: A Knowledge-Based Approach for Interpreting Genome-Wide Expression Profiles.” Proceedings of the National Academy of Sciences of the United States of America 102 (43): 15545–50.

Suel, G. M., R. P. Kulkarni, J. Dworkin, J. Garcia-Ojalvo, and M. B. Elowitz. 2007. “Tunability and Noise Dependence in Differentiation Dynamics.” Science 315 (5819): 1716–19.

Süel, Gürol M., Jordi Garcia-Ojalvo, Louisa M Liberman, and Michael B Elowitz 2006. “An Excitable Gene Regulatory Circuit Induces Transient Cellular Differentiation.” Nature 440 (7083): 545–50.

Trapnell, Cole, Brian A. Williams, Geo Pertea, Ali Mortazavi, Gordon Kwan, Marijke J. van Baren, Steven L. Salzberg, Barbara J. Wold, and Lior Pachter. 2010. “Transcript Assembly and Quantification by RNA-Seq Reveals Unannotated Transcripts and Isoform Switching during Cell Differentiation.” Nature Biotechnology 28 (5): 511–15.

Tu, Benjamin P., Andrzej Kudlicki, Maga Rowicka, and Steven L. McKnight. 2005. “Logic of the Yeast Metabolic Cycle: Temporal Compartmentalization of Cellular Processes.” Science 310 (5751): 1152–58.

Veening, Jan-Willem, Wiep Klaas Smits, and Oscar P. Kuipers. 2008. “Bistability, Epigenetics, and Bet-Hedging in Bacteria.” Annual Review of Microbiology 62: 193–210.

Wang Ping, Lydia Robert, James Pelletier, Wei Lien Dang, Francois Taddei, Andrew Wright, and Suckjoon Jun. 2010. “Robust Growth of Escherichia Coli.” Current Biology: CB 20 (12): 10991103.

Wolfe,Alan J. 2005. “The Acetate Switch.” Microbiology and Molecular Biology Reviews: MMBR 69 (1): 12–50.

Xiao, Zijun, and Ping Xu. 2007. “Acetoin Metabolism in Bacteria.” Critical Reviews in Microbiology 33 (2): 127–40.

Young, Jonathan W., James C. W. Locke, Alphan Altinok, Nitzan Rosenfeld, Tigran Bacarian, Peter S. Swain, Eric Mjolsness, and Michael B. Elowitz. 2011a. “Measuring Single-Cell Gene Expression Dynamics in Bacteria Using Fluorescence Time-Lapse Microscopy.” Nature

